# The burden of dengue fever and chikungunya in southern coastal Ecuador: Epidemiology, clinical presentation, and phylogenetics from the first two years of a prospective study

**DOI:** 10.1101/102004

**Authors:** Anna M. Stewart-Ibarra, Sadie J. Ryan, Aileen Kenneson, Christine A. King, Mark Abbott, Arturo Barbachano-Guerrero, Efraín Beltrán-Ayala, Mercy J. Borbor-Cordova, Washington B. Cárdenas, Cinthya Cueva, Julia L. Finkelstein, Christina D. Lupone, Richard G. Jarman, Irina Maljkovic Berry, Saurabh Mehta, Mark Polhemus, Mercy Silva, Timothy P. Endy

## Abstract

Here we report the findings from the first two years of an arbovirus surveillance study conducted in Machala, Ecuador, a dengue endemic region (2014-2015). Patients with suspected dengue virus (DENV) infections (index cases, n=324) were referred from five Ministry of Health clinical sites. A subset of DENV positive index cases (n = 44) were selected, and individuals from the index household and four neighboring homes within 200-meters were recruited (n = 400). Individuals who entered the study, other than index cases, are referred to as associates. In 2014, 70.9% of index cases and 35.6% of associates had acute or recent DENV infections. In 2015, 28.3% of index cases and 12.8% of associates had acute or recent DENV infections. For every DENV infection captured by passive surveillance, we detected an additional three acute or recent DENV infections in associates. Of associates with acute DENV infections, 68% reported dengue-like symptoms, with the highest prevalence of symptomatic acute infections in children under 10 years of age. The first chikungunya virus (CHIKV) infections were detected on epidemiological week 12 in 2015. 43.1% of index cases and 3.5% of associates had acute CHIKV infections. No Zika virus infections were detected. Phylogenetic analyses of isolates of DENV from 2014 revealed genetic relatedness and shared ancestry of DENV1, DENV2 and DENV4 genomes from Ecuador with those from Venezuela and Colombia, indicating presence of viral flow between Ecuador and surrounding countries. Enhanced surveillance studies, such as this, provide high-resolution data on symptomatic and inapparent infections across the population.

## Introduction

The region of the Americas is facing an unprecedented public health crisis of co-occurring epidemics of illness due to dengue virus (DENV), chikungunya virus (CHIKV) and Zika virus (ZIKV). These arboviruses cause acute febrile illness and are transmitted to humans by the female *Aedes aegypti* and *Ae. albopictus* mosquitoes.

Dengue fever is caused by an infection by one of the serotypes of the mosquito-borne dengue virus (DENV 1-4, family *Flaviviridae*, genus *Flavivirus*). Clinical manifestations range from mild illness (*i.e.*, fever, rash, joint pain) to severe illness characterized by pathologic vascular permeability leading to hemorrhage, shock, and sometimes death.^1^ Over the last three decades, the distribution, severity, and incidence of DENV has increased in Latin America, from 16.4 cases per 100,000 in the 1980’s to 71.5 cases per 100,000 from 2000 to 2007.^2,3^ Current estimates of apparent DENV infection in the Americas range from 1.5 million^4^ to 13.3 million^5^ infections per year. In 2015, 2.35 million DENV infections were reported in the Americas, leading to 10,200 severe infections and 1,181 deaths.^6^

More recently, CHIKV and ZIKV have emerged and caused major epidemics in the same populations in the Americas. The first CHIKV infections (family *Togaviridae*, genus *alphavirus*) were reported in the Americas in 2013, resulting in over 2.5 million suspected and confirmed cases to date.^7^ The first ZIKV infections (family *Flaviviridae*, genus *flavivirus*) were reported in Brazil in 2015.^8,9^ To date, 805,703 suspected and confirmed cases of ZIKV have been reported from the Americas (as of Nov 30, 2017).^10^

In Ecuador, DENV causes the greatest burden of mosquito-borne febrile illness. In 2014 and 2015, the years of this study, 16,908 and 44,104 cases per year, respectively, were reported.^11^ Historically, DENV was eliminated from Ecuador in the 1950s through the use of DDT and other measures to control *Ae. aegypti*, the only known vector in Ecuador.^12,13^ Following a weakening of the vector control program and the re-invasion of *Ae. aegypti* in the 1970s and 1980s, DENV1 re-emerged in Ecuador in 1988, and caused a major epidemic of classic dengue fever.^14^ From 1993 to 1999 three serotypes circulated: DENV1, DENV2 (American strain), and DENV4. In 2000, DENV3 and DENV2 (Asian strain) were identified, and the first cases of severe hemorrhagic dengue were subsequently reported.^15^

Today the burden of DENV is greatest in the coastal lowland region of Ecuador, the site of the current study. Prior studies in southern coastal Ecuador indicate that DENV transmission is highly seasonal, with the greatest incidence of disease and density of mosquito vectors from February to May, the hot and rainy season, and lower transmission throughout the rest of the year.^16,17^ DENV epidemics in the region are associated with El Niño climate events that result in warmer air temperatures.^16^ Local social-ecological risk factors for DENV infections and *Ae. aegypti* proliferation in this region include adjacent abandoned properties, interruptions in piped water, shaded patios, lack of use of mosquito bed nets, lack of fumigation inside the home, poor housing conditions, inadequate piped water, gaps in knowledge about DENV transmission, and water storage habits.^17^^‒^^20^

The first autochthonous CHIKV infections were reported in Ecuador at the end of 2014; to date 35,891 suspected and confirmed cases have been reported (as of Nov 30, 2017).^7^ The first autochthonous ZIKV infections were confirmed in Ecuador on January 7, 2016. A total of 6,240 suspected and confirmed cases of ZIKV have been reported (as of Nov 30, 2017), including seven cases of congenital syndrome associated with ZIKV, which were first reported in May 2017.^10^

In Ecuador, suspected and confirmed DENV, CHIKV, and ZIKV cases require mandatory notification to the Ministry of Health (MoH). The MoH in Ecuador follows the 2009 World Health Organization (WHO) dengue diagnostic guidelines.^1^ The national surveillance system is based on passive surveillance of cases from MoH clinics and hospitals. A subset of suspected cases are confirmed for DENV using nonstructural protein 1 (NS1) antigen and immunoglobulin (IgM) ELISAs in local diagnostic laboratories operated by the MoH. A subset of cases are confirmed for DENV, CHIKV, and ZIKV using quantitative PCR at the national reference laboratory of the National Institute for Public Health Research (INSPI) of the MoH. Suspected infections trigger focal vector control interventions in the infected home and surrounding homes by the MoH (i.e., fogging, indoor residual spraying, source reduction, and larvicide application).

There have been prior enhanced surveillance studies to estimate the burden of dengue fever in Asia^21^^‒^^24^ and Latin America^25^^‒^^31^, with study designs ranging from pediatric to adult cohorts, tracking of school-based absentees, use of sentinel clinics, and community-based cluster investigations. In general, these studies found that enhanced surveillance methods identified a greater number of DENV infections, especially mild and inapparent infections, compared to traditional passive surveillance systems. Enhanced surveillance studies generate high-resolution data on the spatio-temporal distribution of symptomatic and inapparent infections across the population. This is especially important in settings and in subgroups with low-health care seeking behavior or limited access to health centers. These data allow the public health sector to more accurately estimate the social and economic burden of the disease, allowing for more informed decision-making regarding the allocation of scarce resources. These studies can also inform the design and implementation of interventions targeted at high-risk groups, such as vaccination campaigns or vaccine trials.

Here we present the results of the first two years of an active surveillance study in Ecuador. The aim of this study was to characterize the epidemiology, clinical presentation, and viral phylogenetics of DENV. We also present the epidemiology and clinical characteristics of CHIKV during the first CHIKV outbreak. This study is part of a long-term partnership with the MoH of Ecuador focused on strengthening febrile vector-borne disease surveillance in southern coastal Ecuador, providing high resolution epidemiological information for the region.^32^

## Materials and Methods

### Definitions

*Index cases* are hospitalized patients and outpatients with a clinical diagnosis of an acute DENV infection who enrolled in the study. *Initiate index cases* are index cases that tested positive for DENV and were randomly selected to initiate a cluster investigation. *Associates* are study subjects who resided in the home of the initiate index case and/or in the four neighboring homes located in the cardinal directions at a maximum distance of 200 meters from the initiate index household. The four associate homes plus the initiate index case home are referred to as a *cluster*.

A study subject was considered to have an *acute DENV infection* if s/he tested positive by NS1 rapid test, NS1 ELISA or RT-PCR. If the person was negative for those three tests, but was positive by IgM ELISA, they were classified as having a *recent DENV infection*. Individuals were classified as *uninfected with DENV* if they were negative for NS1 rapid test, NS1 ELISA, RT-PCR and IgM ELISA. Individuals who tested negative for all of the tests except for the presence of IgG antibodies were not classified. Individuals who tested positive for CHIKV or ZIKV by RT-PCR were classified as having an *acute CHIKV* or *acute ZIKV infection*.

We define a *symptomatic* individual as an associate with one or more dengue-like symptoms. By definition, all index cases are symptomatic. Prior studies that report symptomatic illness, defined symptomatic as febrile,^24,33^ whereas we use a broader definition of symptomatic to include any dengue-like symptom (*e.g.*, headache, muscle/joint pain, retro-orbital pain, abdominal pain, drowsiness/lethargy, fever, rash), since symptoms other than fever were more frequently reported by associates with acute DENV infections (Supplementary Table 1). An *inapparent* infection is defined as an infection in an associate who has no dengue-like symptoms.

### Ethics Statement

This study protocol was reviewed and approval by Institutional Review Boards (IRBs) at SUNY Upstate Medical University, Cornell University, the Human Research Protection Office (HRPO) of the U.S. Department of Defense, the Luis Vernaza Hospital in Guayaquil, Ecuador, and the Ecuadorean Ministry of Health. Prior to the start of the study, all participants engaged in a written informed consent or assent process, as applicable. If the participant was unable to participate in the consent or assent process, an adult representative documented their consent. Children aged 7 to 17 signed an assent statement and parents signed an informed consent. Parents signed an informed consent on behalf of children under 7 years to > 6 months. The study included children (> 6 months) to adults (index cases) who were evaluated in sentinel clinics or the hospital with a clinical diagnosis of acute DENV infection. Before signing the informed consent, index cases were informed that they might be randomly selected to participate in a cluster investigation (initiate index cases). Additional study subjects include associate children (> 6 months) and adults, who resided in the cluster homes.

### Study Site

Machala, Ecuador, (population 280,694, capital of El Oro Province) is a port city located along the Pan American Highway, near the Ecuador-Peru border (Fig 1). Machala has among the highest incidence rates of DENV in Ecuador and exceptionally high *Ae. aegypti* densities compared to other countries in Latin America and Asia.^17,34,35^ In 2014 and 2015, 1,196 and 2,791 DENV cases, respectively, were reported from Machala (annual incidence of 42.6 cases per 10,000 people in 2014, 99.4 cases per 10,000 people in 2015).^36^ The first local cases of CHIKV were reported by the MoH in May 2015, and the first cases of ZIKV were reported in February 2016. Machala is a strategic location to monitor and investigate DENV -- and now CHIKV and ZIKV -- transmission dynamics due to its location near an international border and port, and the historically high incidence of mosquito-borne diseases.

**Fig 1:**
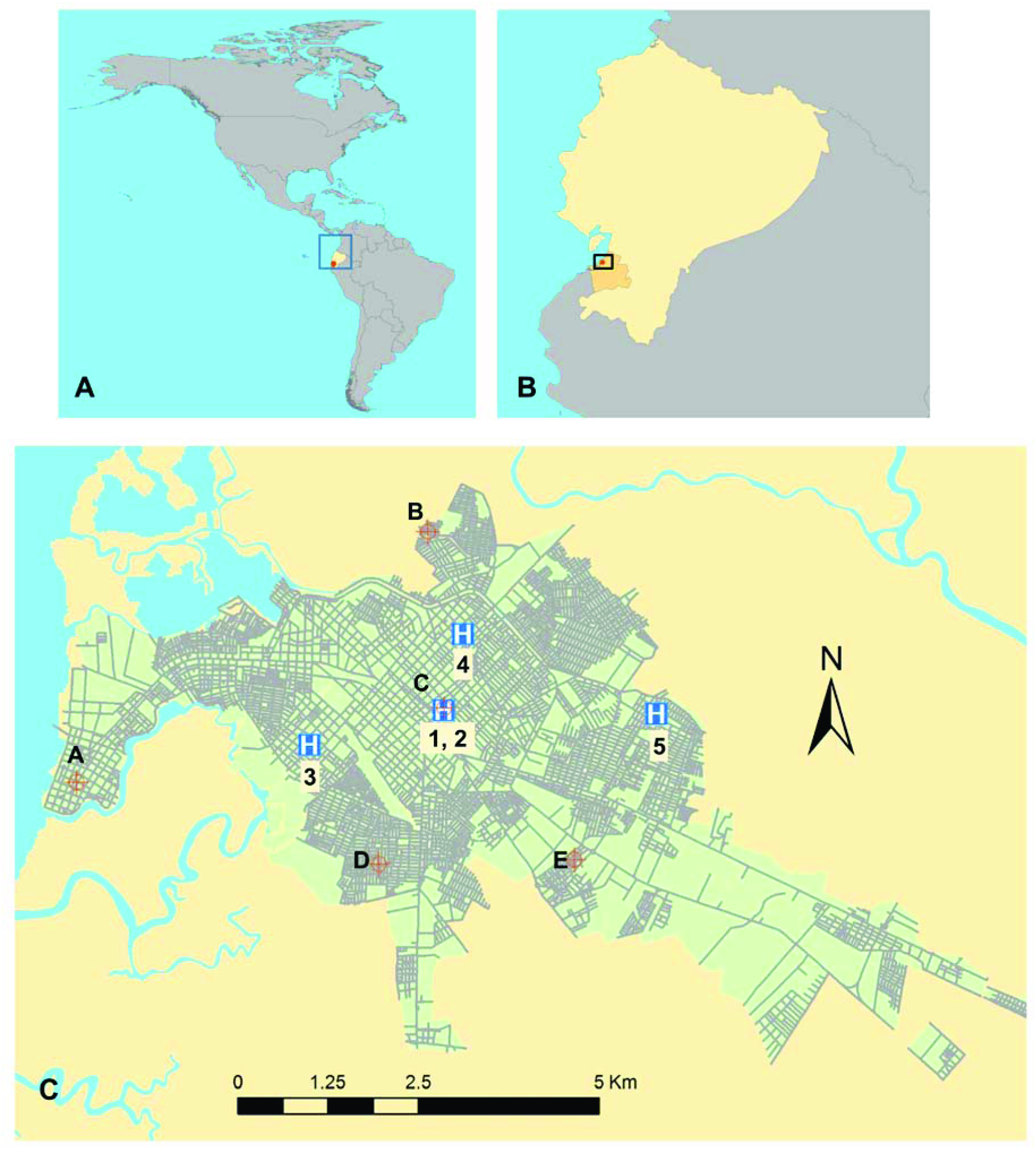
Map of the study site: A. Location of Ecuador in the Americas. B. Location of El Oro Province in Ecuador, the city of Machala indicated as a red dot. C. The city of Machala, showing the five Ministry of Health clinical sites/hospital: 1. Mabel Estupiñan Clinic, 2. Teofilo Davila Hospital, 3. Brisas del Mar Clinic, 4. El Paraiso Clinic, 5. Rayito de Luz Clinic. The location of meteorological stations are indicated by A-E as follows: A. Puerto Bolivar, B. Los Esteros, C. Mabel Estupiñan; D. Florida; E. Crucitas.

Sentinel clinical sites operated by the MoH in Machala were selected based on historical reported DENV cases and the resources that they were able to offer for coordinating and supporting the methods of this surveillance study. Of the twenty-three MoH clinics in Machala, four were selected. These included the clinics Brisas del Mar, Rayito de Luz, Mabel Estupiñan, and El Paraiso. In addition, the Teófilo Dávila Hospital of the MoH was included, because it is the principal public hospital of the province, where the MoH clinics refer patients with severe DENV infections.

### Passive and active surveillance study design

Hospitalized patients and outpatients with a clinical diagnosis of an acute DENV infection (index cases), as determined by MoH physicians, were referred to our study technician or nurse and were invited to participate in the study. Consent was obtained and the following data were collected using a customized database on an Ipad (FileMaker Pro Advanced 13.0v5): patient demographics, home address, primary reason for seeking medical care, date of onset of fever, symptoms within the last seven days, medications, and aural temperature. Data were uploaded daily and stored in a secure cloud-based server (GoZync). At the time of clinical evaluation, a 20 ml blood specimen (adjusted for age and weight by the National Institute of Health criteria) was obtained by venipuncture from each participant. Samples were processed at our diagnostic laboratory at the hospital. Serum samples were used to test for acute DENV infections using NS1 rapid strip tests (PanBio Dengue Early Rapid Test). NS1 tests were run the same day that the index case was recruited into the study. Additional serum, cells, and plasma were separated via centrifugation and aliquoted in multiple tubes and stored at -80**°**C.

Each week, up to four index cases that were positive for DENV infection were randomly selected to be initiate index cases, and they were invited to participate in the active surveillance component of this study. The study team visited the household of the initiate index case and the nearest neighboring homes in each of the four cardinal directions, at a distance of less than 200 meters from the index household, the typical flight range of the *Ae. aegypti* mosquito. All household members (associates) from this cluster of homes were invited to participate in the study. Investigations in clusters began within two days of the initiate index case entering the study. The diagnostic tests and clinical assessments described above for index cases were repeated for all associates. The location (latitude, longitude) of each home was recorded using handheld Garmin GPS units. Passive and active surveillance study designs were optimized in a prior study by the Armed Forces Research Institute of Medical Sciences (AFRIMS) in Kamphaeng Phet Province, Thailand.^24^

### Diagnostic assays

Additional diagnostic testing for DENV was conducted using serum samples and commercial ELISA kits (Panbio) to test for NS1 (Dengue Early ELISA), IgM (Dengue Capture IgM), and IgG (Dengue Capture IgG). We classified participants as having a primary DENV infection if the ratio of IgM to IgG was ≥ 1.8, and a secondary DENV infection if the ratio was less than 1.8.^24,37,38^

Specimens were shipped to SUNY Upstate Medical University for testing by qualitative real-time reverse transcriptase (RT)-PCR assays for DENV1-4, CHIKV, and ZIKV. All samples from 2014 and 2015 were screened for DENV1-4. Samples from index cases in 2014 and index cases and associates in 2015 were screened for CHIKV. Only samples from index cases and associate in 2015 were screened for ZIKV. All analyses were performed on a BioRad DNA Engine Chromo 4 System with MJ Opticon Monitor Analysis Software. For DENV1-4 analysis, total RNA was extracted from 140 µL of human serum specimens using the QIAamp® Viral RNA Mini Kit (QIAgen, Cat# 52906) according to the manufacturer’s suggested protocol and resuspended in 50 µL of buffer. Ten (10) µL of RNA (or the equivalent of 28 µL of serum) was used in a 20 µL reverse transcriptase reaction, of which 5 µL of the resulting cDNA was used for the PCR reaction. All samples and controls were analyzed in duplicate in a multiplex RT-PCR reaction for 45 cycles using SuperScript III Platinum One-Step qRT-PCR System (Life Technologies Cat# 11732-020) based on the CDC DENV1-4 Real Time RT-PCR Assay (CDC, Catalog number KK0128)^39^ and a published assay.^40^ Samples were classified as positive according to a suggested C(t) value of ≤ 37.0, which coincides with a cutoff based on CDC recommendations for identifying positive DENV samples.^39^ For ZIKV and CHIKV analysis, total RNA was extracted from human serum specimens using the QIAamp® Viral RNA Mini Kit (QIAgen, Cat# 52906) according to a modified assay developed at the Walter Reed Army Institute of Research (WRAIR), Viral Diseases Branch. All samples and controls were analyzed in duplicate in a multiplex RT-PCR reaction using TAQMAN Fast Virus 1-Step Mix (Life Technologies Cat# 4444432). The CHIKV primer and probe set (HEX reporter) was adapted from an AFRIMS protocol, Set 3, which was designed specifically for the Asian genotype CHIKV strain currently in the Caribbean and verified using Synthetic CHIKV RNA control (ATCC, Cat# VR-3246SD). The ZIKV primer and probe set (FAM reporter) was based on the AFRIMS protocol that was adapted from a published assay^41^ and verified using RNA extracted from ZIKV culture fluid (ZeptoMetrix Corp., Cat# 0810092CF). Both primer/probe sets were specific for their respective viral target and did not detect other viruses (DENV1-4, YFV, and JEV). Samples were classified as positive based on the same cutoff value used for DENV (C(t) value of ≤37.0). Primers and probes for DENV, CHIKV, and ZIKV are shown in Supplementary Table 2.

### Statistical analysis

Statistical analyses were conducted using R (version 3.3.3) in RStudio (version 1.0.136), using the ‘base’ and ‘psych’ packages for summary statistics. Student’s t-test was used to determine differences in continuous variables, and Chi-square or Fisher’s exact test were used for proportions.

### Sequencing and consensus assembly

Samples from 2014 that were DENV positive by RT-PCR were sent to WRAIR, Viral Diseases Branch, for full-length sequencing. Samples were extracted using a QIAGEN QIAamp viral mini RNA extraction kit in accordance with manufacturer’s protocols. Full genome was amplified on Fluidigm Access Array system using DENV serotype specific primers and the Life Technologies SuperScript TM III One-Step RT-PCR system with Platnimum® Taq High Fidelity polymerase, followed by cDNA quality check using Agilent Bioanalyzer DNA7500 kit and RT-PCR product purification. Purified RT-PCR products were quantified using the Invitrogen Quant-iTTM PicoGreen dsDNA Reagent and Kit following the manufacturer’s protocols. MiSeq library preparation included: dilution of purified amplicons products to 0.2ng/µL, tagmentation using 5 microliters of each dilution stock as input DNA, neutralization of each Nextera® XT Tagmentation reaction using 5µl NT buffer, PCR amplification using index primers from Nextera XT Index kit version 2 set C, PCR clean up using 25 microliters per PCR reaction of Beckman Counter AMPure XP beads, and library normalization using applicable reagents provided in the Nextera XT® DNA Library Preparation kit. After normalization, each library was pooled and sequenced using the Illumina MiSeq reagent kit (version 2, 500 cycles) and Illumina MiSeq next generation sequencer in accordance with Illumina protocols.

Construction of consensus genomes was performed using ngs_mapper v1.2.4 in-house developed pipeline (available on github, http://github.com/VBDWRAIR/ngs_mapper). Briefly, raw fastq data were stripped of barcodes and adapters and subjected to read filtering using a quality threshold of Q25. Remaining reads were further end-trimmed using a quality threshold of Q25 using Trimmomatic.^42^ Trimmed reads with quality >Q25 were initially mapped to a set of reference sequences to determine the best reference fit for each of the samples. Following reference determination, reads from each of the samples were re-mapped to their closest related reference genome, to maximize the number of mapped reads. Reference mapping was performed using the BWA-MEM algorithm.^43^ Assemblies were further processed using samtools version 0.1^44^ and an in-house developed python program called *basecaller.py* to produce an adapted VCF for each segment, in parallel, which incorporates genomic ambiguity inherent in RNA viruses into the final consensus genome for that sample based on thresholds set by the investigator. Threshold for consensus genomic reconstruction for ambiguity incorporation was set at 20% for this analysis, meaning if any site contained a different nucleotide call that was present at 20% or greater in the dataset (taking quality of call into account) the site was given an ambiguous base call (according to IUPAC conventions). Consensus sequences for all samples were constructed, in parallel, from the adapted VCF output. All consensus sequences were further manually quality-checked. Statistics and graphics illustrating read depth and quality of mappings for each sample across each segment produced by the pipeline were done using matplotlib.^45^

### Phylogenetic analyses

The five sequenced full genome DENV1 samples were aligned to a set of full genome DENV1 reference sequences obtained from GenBank using MEGAv6.^46^ The 131 reference genomes were selected to represent: i) all DENV1 genotype lineages, for accurate genotype determination, ii) wide sampling time periods, with a focus on the most recently sampled genomes (2009-2016), iii) most geographical regions, with a focus on Central and South America. In addition, the top 20 genomes matching the five genomes from Ecuador through Basic Local Alignment Search Tool (Blast)^47^ were added to the reference dataset. A set of 140 full genome DENV2 reference sequences was obtained from GenBank following the same criteria as for DENV1, and aligned to the 27 DENV2 sequenced genomes from Ecuador. Likewise, a set of 100 full genome DENV4 reference sequences was obtained from GenBank following the same criteria as for DENV1, and aligned to the single DENV4 sequenced genome from Ecuador. We were unable to sequence DENV3 due to limited sample volume. Genetic sequences are deposited in GenBank under accession numbers KY474303-KY474335.

We determined the best-fit models of evolution for DENV1, DENV2 and DENV4 datasets using jModelTest v2.1.7 with Akaike Information Criterion (AIC) and Bayesian Information Criterion (BIC).^48^ Maximum Likelihood (ML) phylogenetic trees for DENV1, DENV2 and DENV4 datasets were inferred using Phyml v 4.9.1.^49,50^ The model of evolution used for the full genome tree inferences was GTR+I+Γ (general time reversible with empirically estimated proportion of invariant sites and gamma distribution of among-site variation, 4 categories), for all three DENV serotypes. The tree space was searched heuristically using the best of NNI (Nearest Neighbor Interchanges) and SPR (Subtree Pruning and Regrafting). Node confidence values were determined by aLRT (approximate Likelihood Ratio Test) using the nonparametric Shimodaira-Hasegawa approach. Node confidence values of >0.75 are considered good support. The resulting trees were rooted by the KR919820 sylvatic reference genome^51^ for DENV1, and by the sylvatic genotype outgroups for DENV2 and DENV4.

## Results

From January 1, 2014, through December 31, 2015, we recruited 324 index cases with suspected DENV infections from the five clinical sites in Machala, Ecuador (Figs 1 and 2). A subset of 310 index cases (186 in 2014, 124 in 2015) had valid test results and were included in this study (Table 1). A total of 72 index cases were positive by NS1 rapid test, and from these we randomly selected 44 initiate index cases, from which 400 associates were recruited into the study. A subset of 384 associates (298 in 2014, 86 in 2015) had valid test results and were included in this study.

**Fig 2.**
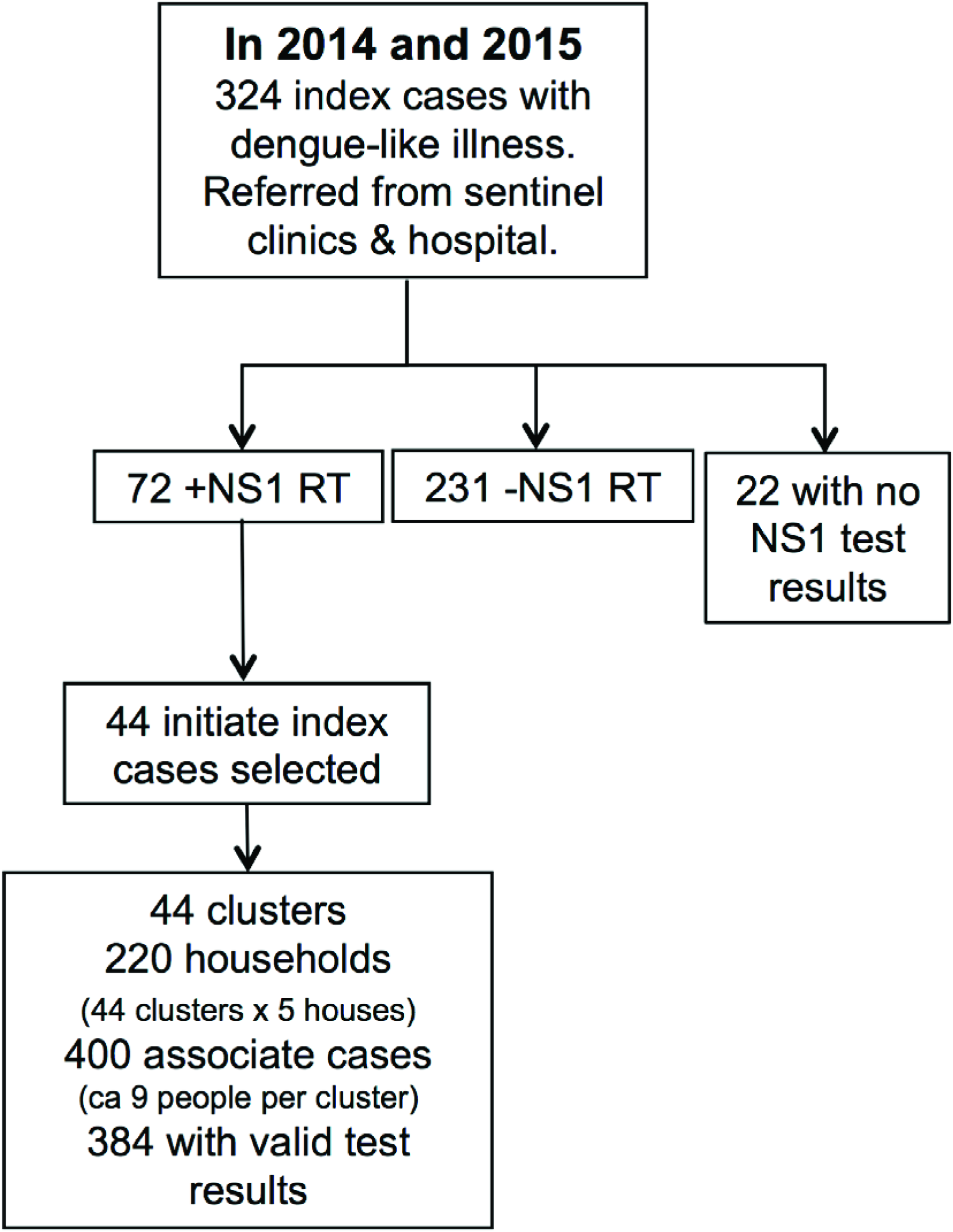
Study design. DENV surveillance study design in Machala, Ecuador.

**Table 1.** Demographic data and infection status of index cases and associates. The characteristics of index cases and associates in 2014 and 2015: mean age (standard deviation = SD) and gender, febrile status, hospitalization status, and arbovirus infection status (DENV acute infection: NS1 RT, NS1 ELISA or RT-PCR positive; DENV recent infection: IgM positive and NS1 RT/NS1 ELISA/RT-PCR negative; CHIKV and ZIKV confirmed by RT-PCR).

|  | 2014 |  | 2015 |  |
| --- | --- | --- | --- | --- |
|  | Index cases | Associates | Index cases | Associates |
|  | N = 186 | N = 298 | N = 124 | N = 86 |
| Age in years, mean (SD) | 20.6 (15.5) | 35.3 (19.1) | 28.0 (18.6) | 38.8 (20.0) |
| Gender, % female | 90/186 (48.4%) | 195/295 (66.1%) | 68/124 (54.8%) | 58/86 (67.4%) |
| Temperature > 38°C | 30/185 (16.2%) | 2/290 (0.7%) | 23/124 (18.5%) | 0/86 (0%) |
| Fever in the prior 7 days | 179/185 (96.8%) | 33/285 (11.6%) | 119/124 (96.0%) | 3/83 (3.6%) |
| DENV infection |  |  |  |  |
| Acute infection | 75/186 (40.3%) | 45/298 (15.1%) | 24/124 (19.4%) | 5/86 (5.8%) |
| Recent infection | 57/186 (30.6%) | 61/298 (20.5%) | 11/124 (8.9%) | 6/86 (7.0%) |
| Hospitalized | 34/186 (18.3%) | Not applicable | 21/124 (16.9%) | Not applicable |
| Other acute infections |  |  |  |  |
| Chikungunya virus | 0/152 (0%) | Not applicable | 53/123 (43.1%) | 3/86 (3.5%) |
| Zika virus | Not applicable | Not applicable | 0/123 (0%) | 0/86 (0%) |

DENV transmission was highly seasonal in 2014 and 2015, with a peak in May (Fig 3). CHIKV was first identified in our study on epidemiological week 12 in 2015, and transmission followed a similar seasonal curve as DENV (Fig 3). No ZIKV infections were detected (Table 1).

**Fig 3.**
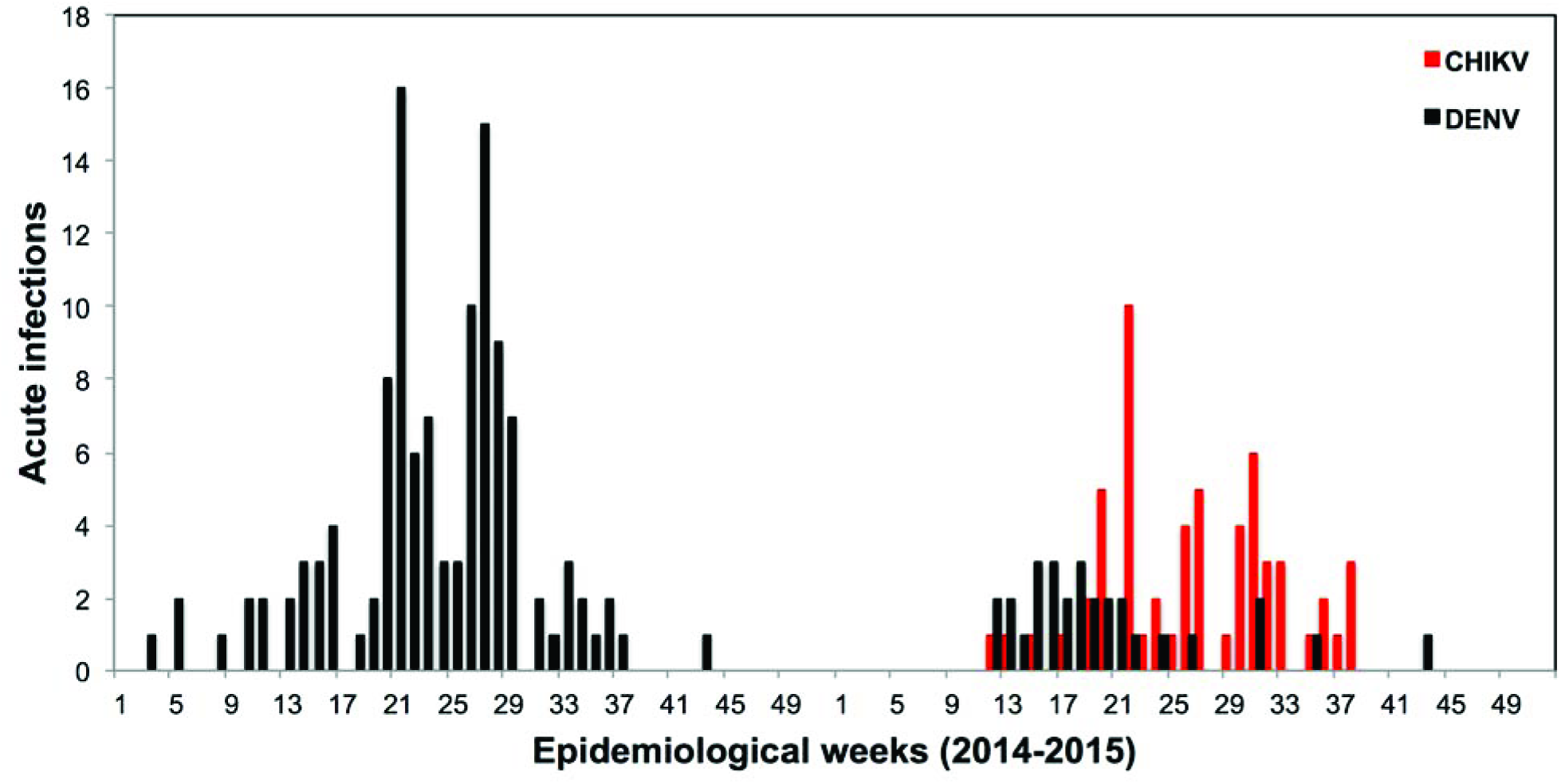
Weekly laboratory confirmed acute DENV and CHIKV infections in 2014 and 2015 detected by passive and active surveillance. Note: no surveillance was conducted in week 30 of 2014.

Table 1 shows the diagnostic results from 2014 and 2015. There were some individuals who did not have enough information to categorize as DENV positive or negative, for example, an individual who was negative for an NS1 rapid test and PCR, but did not have any ELISA or serology test results. To account for these discrepancies, prevalence estimates include people for whom test results were available, as indicated by the denominators in the diagnostic results section of the table.

### Passive surveillance of index cases

In 2014, the majority of all index cases (132/186, 70.9%) were positive for an acute or recent DENV infection (Table 1). All four DENV serotypes were detected, and DENV2 was the predominant serotype (43/51, 84.3% of serotyped index cases) (Table 2). One individual was positive for DENV1 and DENV2. Secondary DENV infections were most prevalent (73/99, 73.7% of index cases with serology and acute or recent DENV infections) (Table 3). Index cases with acute DENV infections were on average 20.7 years of age (SD=15.7) and 62.7% were male (Table 4). The majority reported a fever within the last seven days (97.3%), 21.3% had fever (>38**°**C) upon entering the study, and 16.0% were hospitalized.

**Table 2.** DENV serotypes. Results from the analysis of samples from 69 individuals in 2014 and 24 individuals in 2015 that were serotyped for DENV by RT-PCR. In 2014, all four DENV serotypes were detected, with DENV2 as the predominant serotype. One index case in 2014 was positive for DENV1 and DENV2. In 2015, DENV1 and DENV2 co-circulated, and DENV1 was the predominant serotype.

| DENV<br>serotypes | 2014 |  | 2015 |  |
| --- | --- | --- | --- | --- |
|  | Index cases<br>N = 51 | Associates<br>N = 18 | Index cases<br>N = 23 | Associates<br>N = 1 |
| 1 | 4/51 (7.8%) | 3/18 (16.7%) | 14/23 (60.9%) | 0/1 (0%) |
| 1 & 2 | 1/51 (2.0%) | 0/18 (0%) | 0/23 (0%) | 0/1 (0%) |
| 2 | 43/51 (84.3%) | 10/18 (55.6%) | 9/23 (39.1%) | 1/1 (100%) |
| 3 | 2/51 (3.9%) | 5/18 (27.8%) | 0/23 (0%) | 0/1 (0%) |
| 4 | 1/51 (2.0%) | 0/18 (0%) | 0/23 (0%) | 0/1 (0%) |

**Table 3.** DENV serology results for index cases and associates. The prevalence of primary and secondary DENV infections as a proportion of individuals who had an acute or recent DENV infection and had valid serology results (217/284 individuals with acute or recent DENV infections, as reported in Table 1). Secondary DENV infections were more prevalent in 2014, whereas primary DENV infections were more prevalent in 2015. The serology of index cases in 2014 versus 2015 was significantly different (p<0.001). The serology of associates in 2014 versus 2015 was not significantly different (p>0.05).

| Serology | 2014 |  | 2015 |  |
| --- | --- | --- | --- | --- |
|  | Index cases | Associates | Index cases | Associates |
|  | N = 99 | N = 81 | N = 31 | N = 6 |
| Primary DENV infection | 26 (26.3%) | 38 (46.9%) | 21 (67.7%)* | 4 (66.7%)* |
| Secondary DENV infection | 73 (73.7%) | 43 (53.0%) | 10 (32.2%)** | 2 (33.3%) |

\*Includes 4 index cases and 1 associate with acute CHIKV infections
\*\*Includes 1 index cases with acute CHIKV infections

*Includes 4 index cases and 1 associate with acute CHIKV infections

**Includes 1 index cases with acute CHIKV infections

**Table 4.** Characteristics of acute DENV infections. Index cases and associates with acute DENV infections in 2014 and 2015: mean age (standard deviation = SD) and gender, febrile status, and the proportion who were hospitalized. There were no significant differences between years (p>0.05).

|  | 2014 |  | 2015 |  |
| --- | --- | --- | --- | --- |
| Characteristics | Index cases | Associates | Index cases | Associates |
|  | N = 75 | N = 45 | N = 24 | N = 5 |
| Age in years, mean (SD) | 20.7 (15.7) | 25.2 (18.6) | 19.3 (12.8) | 19.6 (14.6) |
| Gender, % female | 28/75 (37.3%) | 29/45 (64.4%) | 13/24 (54.1%) | 2/4 (50.0%) |
| Temperature > 38°C | 16/75 (21.3%) | 2/43 (4.7%) | 10/24 (41.7%) | 0/5 (0%) |
| Fever in the last 7 days | 73/75 (97.3%) | 10/41 (24.4%) | 24/24 (100%) | 1/5 (20.0%) |
| Hospitalized | 12/75 (16.0%) | Not applicable | 8/24 (33.3%) | Not applicable |

In 2015, more index cases were positive for acute CHIKV infections (52/123, 43.1%) than for acute or recent DENV infections (35/124, 28.3%). One index case was positive for both acute DENV and CHIKV infections, and five index cases were positive for recent DENV and acute CHIKV infections, resulting in 11.5% (6/52) of CHIKV infections with acute or recent DENV infections. DENV1 was the predominant serotype (14/23, 60.9% of serotyped index cases) (Table 2). Significantly more primary DENV infections were reported in 2015 than in 2014 (21/31, 67.7% of index cases with serology and acute or recent DENV infections, p<0.001, Table 3). Index cases with acute DENV infections were on average 19.3 years of age (SD=12.8), and 54.1% were female (Table 4). All index cases with acute DENV infections reported a fever within the last seven days, 41.7% had fever upon entering the study, and 33.3% were hospitalized. There were no significant differences in the demographics, febrile symptoms, or hospitalization rates for index cases with acute DENV infections between 2014 and 2015 (Table 4, p>0.05).

We estimated the prevalence of symptomatic acute (SA) infections for DENV and CHIKV by age class as a proportion of the total number of individuals recruited per age class (Fig 4, see Supplementary Table 3 for prevalence calculations). Index children 10 to 19 years of age had the highest prevalence of SA DENV infections (40/97, 41.2%). SA DENV prevalence generally declined with increasing age, with the exception of individuals 50 to 59 years of age (7/21, 33.3%). Interestingly, the proportion of primary DENV infections decreased from 0 to 49 years, and increased from 50 to 79 years (as determined by index cases with serology and acute or recent DENV infections). In contrast, the prevalence of SA CHIKV infections, as a proportion of all individuals recruited into the study, was greatest in index cases 60 to 79 years of age (7/9, 77.8%), and prevalence increased with increasing age.

**Fig 4.**
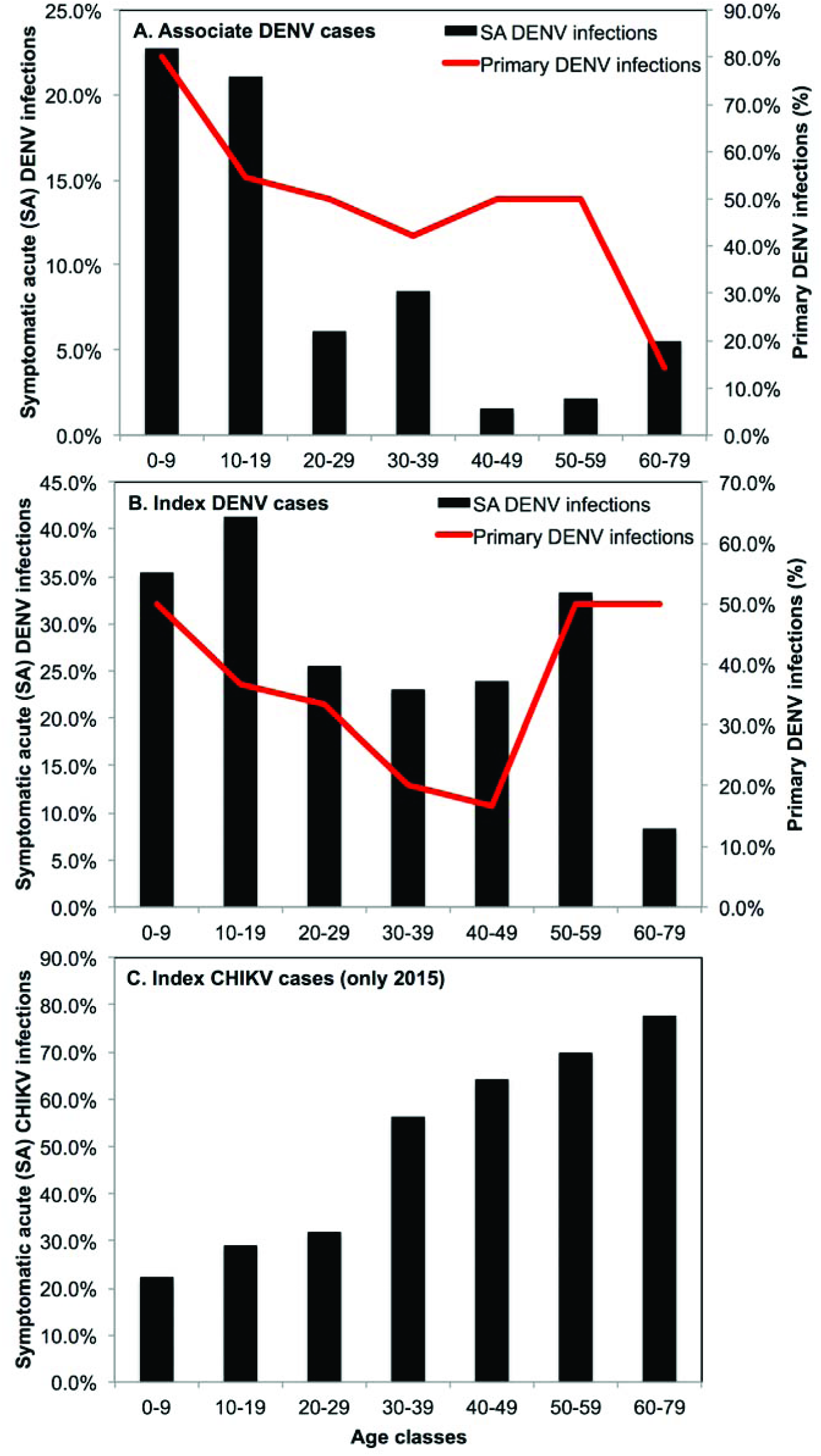
The prevalence of symptomatic acute (SA) infections and serology by age class. The prevalence of SA DENV infections and the proportion of primary DENV infection in 2014 and 2015 for (A) associates and (B) index cases, and (C) the prevalence of SA CHIKV infections in index cases in 2015. See Supplementary Table 3 for raw data and calculation details.

We compared the demographics and symptoms of index cases with acute DENV versus CHIKV infections. Index cases with acute DENV infections were significantly younger (mean=20.2 years, SD=15.0) and more likely to report anorexia and nausea, vomiting and abdominal pain (p<0.05). Index cases with CHIKV were more likely to be female, were older (mean=35.8 years, SD=19.4), and more likely to report muscle or joint pain (p<0.05). A greater proportion of individuals with CHIKV reported rash (CHIKV: 34.6%; DENV: 16.5%; p=0.05), and a lower proportion had fever (> 38°C) upon entering the study (CHIKV: 11.8%, DENV: 26.5%; p=0.06); however, these differences were not statistically significant.

We also compared the demographics and symptoms of primary versus secondary DENV infections (Supplementary Table 4), and DENV1 versus DENV2 infections in index cases (Supplementary Table 5). Individuals with secondary DENV infections were significantly older (secondary: mean=23.2 years, SD=13.8; primary: mean=18.0 years, SD=13.1) (p<0.05). Overall, we identified more severe illness in secondary DENV infections; individuals with secondary infections were more likely to report vomiting, and hospitalized individuals were more likely to have secondary DENV infections (p<0.05). However, individuals with primary DENV infections were more likely to report fever (p<0.05). We did not find significant differences in symptoms between DENV1 and DENV 2 (p>0.05), the predominant serotypes detected in this study, although index cases with DENV2 infections were significantly older (DENV1: mean=14.7 years, SD=10.5; DENV2: mean=25.2 years, SD=16.2) (p<0.05).

### Active surveillance of associates

In each cluster of homes, approximately nine associates were recruited into this study per initiate index case (Fig 2). The distance between the households of associates and the respective initiate index households ranged from 2.2 to 164 meters, with an average of 39 meters (SD=29 m). Most associate households (95.4%) were within 100 meters of the initiate index household.

In 2014, approximately one third of all associates (106/298, 35.6%) had evidence of acute or recent DENV infections (Table 1). As with index cases, DENV2 was the dominant serotype (Table 2). A similar proportion of primary (46.9%) and secondary infections (53.0%) were detected (as determined by associates with serology and acute or recent DENV infections) (Table 3). In 2015, as with index cases, the prevalence of DENV infections decreased as a proportion of all associates recruited (11/86, 12.9%), and primary DENV infections were more common (4/6, 66.7% of associates with serology and acute or recent DENV infections, Table 3). Only one associate was serotyped as DENV2 (Table 2). The serology of associates in 2014 versus 2015 was not significantly different due, in part, to the small sample size (p>0.05). In 2015 we detected acute CHIKV infections in three associates (3/86, 3.5%), including one associate with both acute CHIKV and recent DENV infections.

Approximately two thirds of associates with acute DENV infections (34/50, 68%) reported one or more dengue-like symptoms within the last seven days, resulting in a ratio of symptomatic:inapparent infections (S:I) of 1:0.47 (2.13) (Supplementary Table 1). The most commonly reported symptoms were headache (32%), drowsiness/lethargy (24%), fever (22%), muscle/joint pain (22%), and retro-orbital pain (22%). Only two associates with symptomatic acute DENV infections had sought medical care within the last seven days (2/34, 5.9%), and no associates were hospitalized due to a DENV infection (Table 4). There were no significant differences in the demographics or febrile symptoms of associates with acute DENV infections in 2014 versus 2015 (p>0.05, Table 4).

In associates, we determined the prevalence of SA DENV infections by age class as a proportion of the total number of associates recruited per age class (Fig 4, Supplementary Table 3). Children 0 to 9 years of age had the highest prevalence of SA DENV infections (5/22, 22.7%), and prevalence declined with increasing age. The proportion of primary DENV infections similarly decreased with increasing age. We calculated the prevalence of symptomatic infections in associates with positive primary and secondary DENV infections, and found that individuals with secondary infections had a higher prevalence of symptomatic disease; however, the differences were not statistically significant (symptomatic primary: 24/42, 57.1%; symptomatic secondary 35/45, 77.8%; p=0.07). No associates had SA CHIKV infections.

At the cluster level, prevalence rates varied by the DENV serotype of the initiate index case. In 10 of 44 clusters, the initiate index case had a DENV1 infection. In these clusters, 20% of all associates had acute or recent DENV infections (12/60; 95% CI: 11.8-31.8%), with a range of 0% to 57.1%. The initiate index case had a DENV2 infection in 17 of 44 clusters. Among these clusters, a significantly greater proportion of all associates (36.6%; 59/161; 95% CI: 29.6-44.3%) (p=0.02) had an acute or recent DENV infections, with a range of 12.5% to 87.5%.

We calculated the average number of acute and recent (AR) DENV infections and symptomatic acute and recent (SAR) infections per cluster (see raw data in Supplementary Table 6). By definition, each cluster included an initiate index case, which was a SAR infection. In 2014, there were 32 clusters, with an average of 10.3 (SD=2.7) individuals enrolled per cluster. We detected an average of 4.3 (SD=2.3) AR infections, of which 3.3 (SD=1.7) were SAR infections per cluster. In 2015, there were 12 clusters, with an average of 8.2 (SD=2.2) individuals enrolled per cluster. We detected an average of 1.9 (SD=0.7) AR infections, of which 1.4 (SD=0.7) were SAR infections. All measures were significantly greater in 2014 than in 2015 (p<0.05). Over both years, we detected an average of 3.7 (SD=2.3) AR infections and 2.8 (SD=1.7) SAR infections per cluster.

### Phylogenetic analysis of DENV

The best-fit models for the evolution of DENV1, DENV2, and DENV4, as determined by AIC versus BIC, agreed in all instances. ML phylogenetic tree demonstrated a clear distinction of DENV1 genotypes *I, II, IV* and *V*, and the sylvatic genotypes *III* and *VI* (Fig 5). The five genomes from Ecuador, all sampled in 2014, belonged to genotype *V* of DENV1 and were found in the sub-lineage containing mainly Central and South American genomes (*i.e.*, Colombia, Venezuela, Argentina, Brazil and Puerto Rico). More importantly, sequences from Ecuador fell into two distinct clades within this sub-lineage; two Ecuadorian genomes were more closely related to genomes sampled in Argentina and Venezuela (Clade A), and three Ecuadorian genomes were more closely related to a genome from Colombia (Clade B).

**Fig 5.**
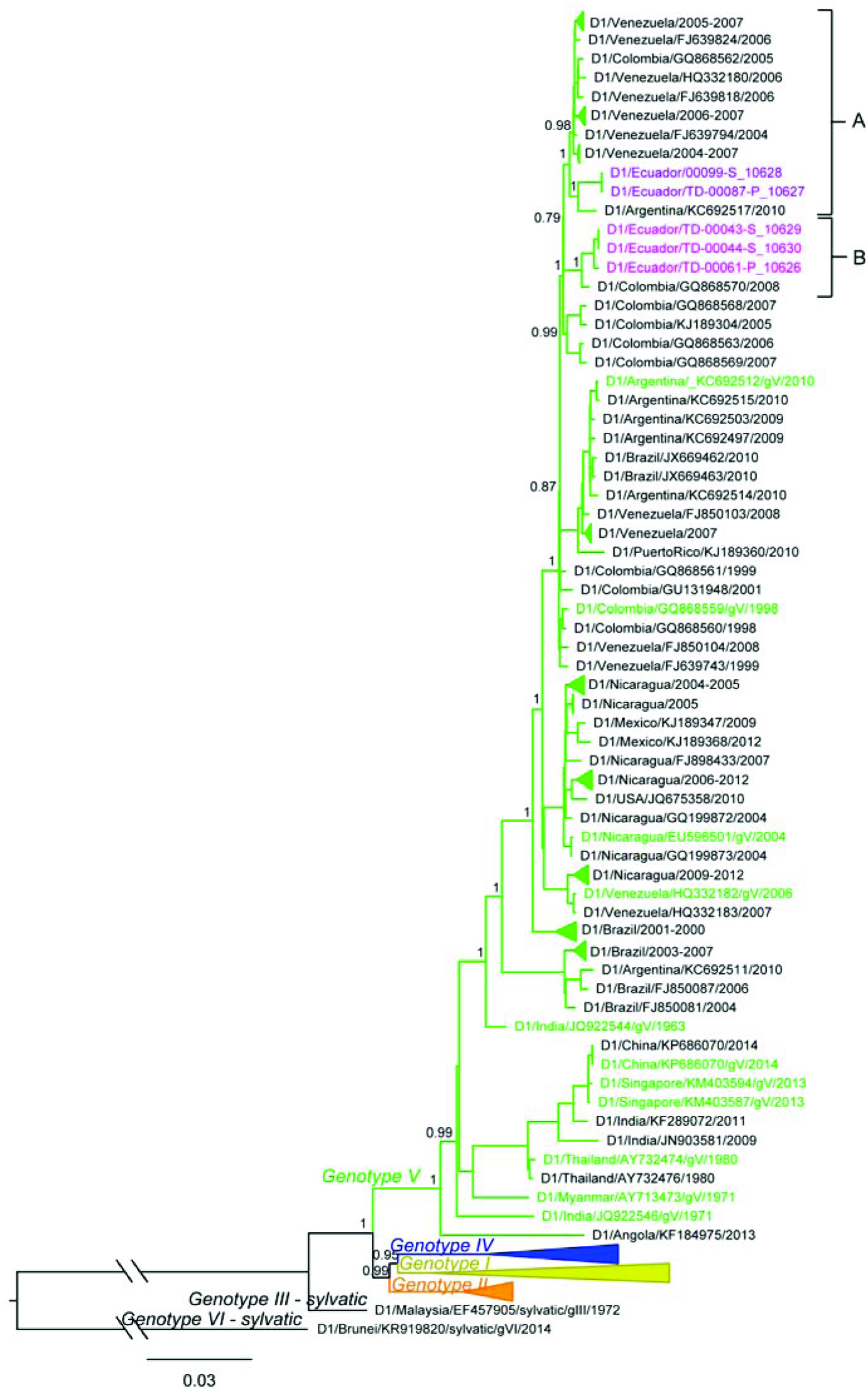
Maximum likelihood phylogenetic tree of DENV1 genotypes from Ecuador in 2014. Samples from Ecuador are colored magenta (dark and light). The two clades containing the genomes from Ecuador are marked in the tree (A and B). aLRT confidence values are shown next to the respective node. The tree is rooted on the sylvatic genotype *VI* sample. Some clades were collapsed in the tree to increase clarity. All collapsed clades were supported with high (>0.75) aLRT values and contained only genomes from a single country, indicated in the name of the clade. Colored taxa represent known genotype references.

The ML phylogenetic tree of DENV2 showed a clear distinction of DENV2 genotypes, including sylvatic, American, Cosmopolitan, Asian I, Asian II and Asian/American (Fig 6). The samples from Ecuador were found within the Asian/American genotype, making up a monophyletic cluster (Clade A) separated from the rest of the South American taxa with high support (aLRT = 1). Genomes clustering closest to the clade A from Ecuador were sampled in Colombia and Venezuela. Sequences from other neighboring countries, such as Peru and Brazil, were found further down in the Asian/American lineage and were separated from the clade A, and from sequences from Colombia and Venezuela, with high support (aLRT = 0.99).

**Fig 6.**
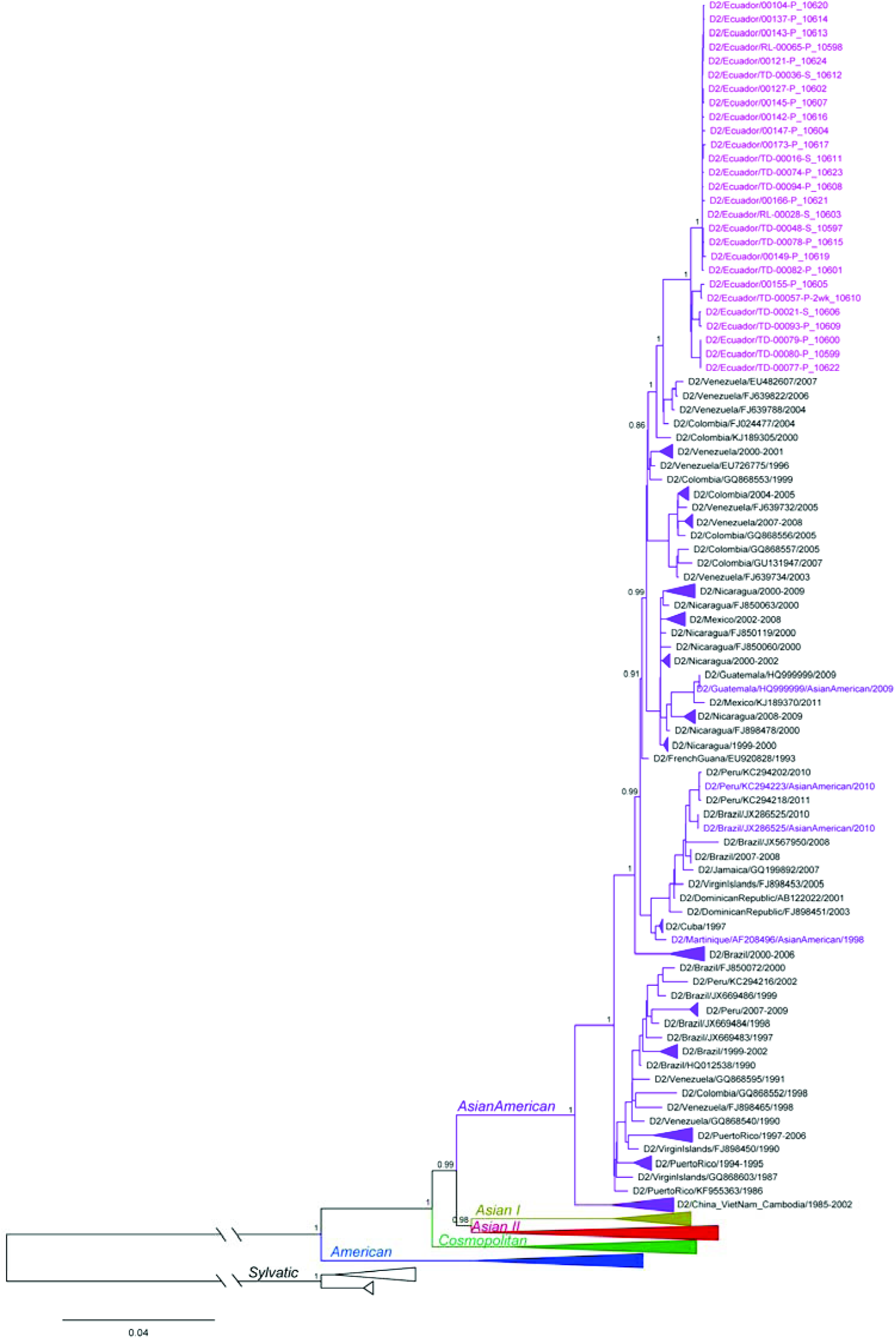
Maximum likelihood phylogenetic tree of DENV2 genotypes from Ecuador in 2014. Samples from Ecuador are colored magenta in a monophyletic clade A. aLRT confidence values are shown next to the respective node. The tree is rooted on the sylvatic genotype outgroup. Some clades were collapsed in the tree to increase clarity. All collapsed clades were supported with high (>0.75) aLRT values and contained only genomes from a single country, indicated in the name of the clade. Colored taxa represent known genotype references.

The ML phylogenetic tree of DENV4 demonstrated a clear distinction of genotypes *I, IIA, IIB, III* and sylvatic (Fig 7). However, two taxa from India/1961-1962 clustered with genotype *I* with low support (aLRT=0.04), indicating that their position in the tree was uncertain and they might belong to a different genotype. The single Ecuador sequence was located within the genotype *IIB* lineage (magenta in the tree). It was surrounded by sequences collected from Venezuela, Colombia and Brazil, indicating their common ancestry. However, the aLRT support for the Ecuador node was low (0.4), suggesting that its correct placement was uncertain.

**Fig 7.**
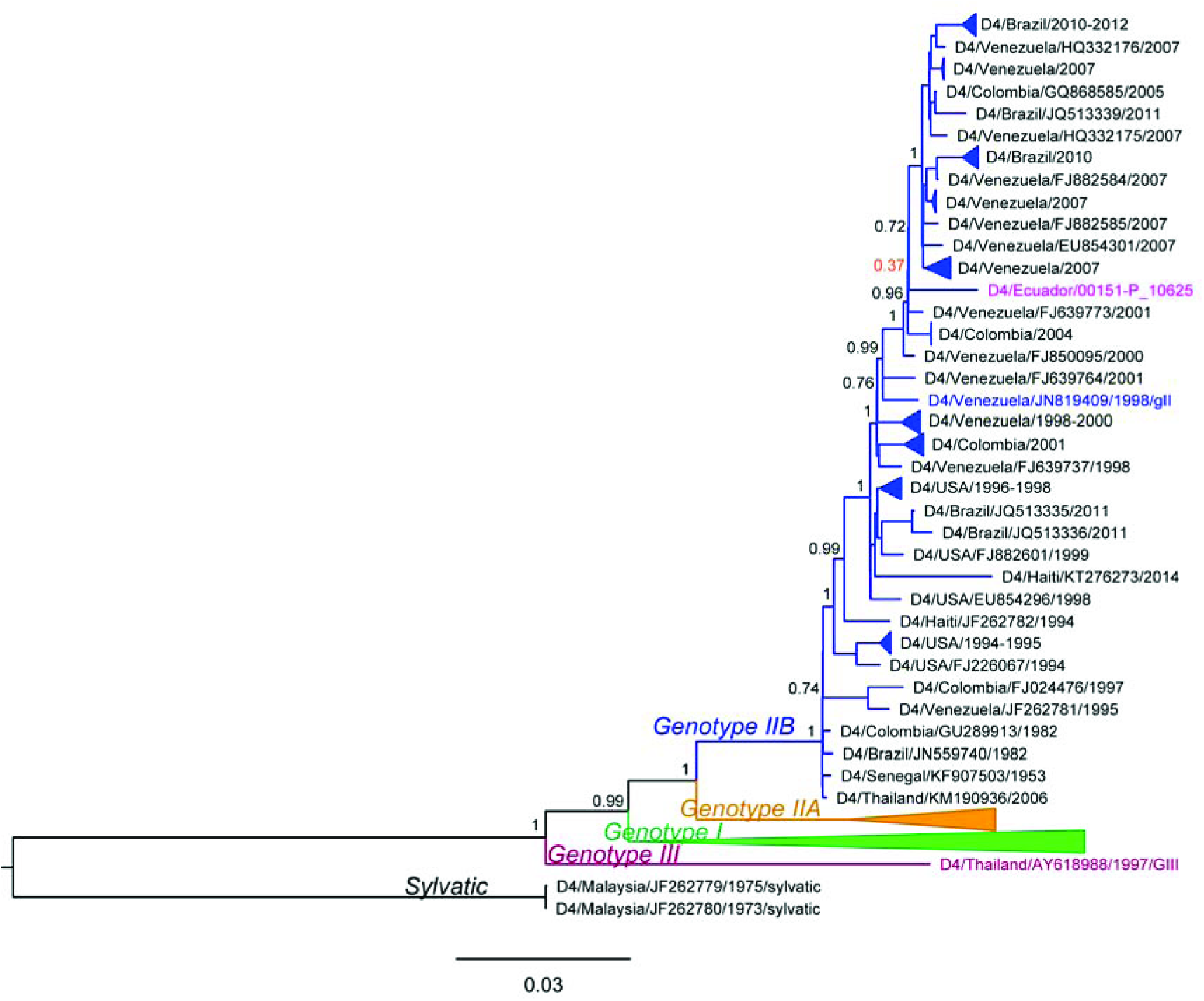
Maximum likelihood phylogenetic tree of DENV4 genotypes from Ecuador in 2014. Sample from Ecuador is colored in magenta. aLRT confidence values are shown next to the respective node. Low aLRT values are highlighted in red. The tree is rooted on the sylvatic genotype outgroup. Some clades were collapsed in the tree to increase clarity. All collapsed clades were supported with high (>0.75) aLRT values and contained only genomes from a single country, indicated in the name of the clade. Colored taxa represent known genotype references.

## Discussion

In this study, we characterized the epidemiology and clinical characteristics of DENV and CHIKV infections, and the phylogenetics of DENV, through an enhanced surveillance study design in an endemic region. We found that burden of symptomatic acute DENV in associates was greatest in children under 10 years of age. In 2014, for every symptomatic acute DENV infection detected by passive surveillance (initiate index cases), we detected an additional three acute or recent infections in associates by active surveillance. Two thirds of associates with acute DENV infections presented with dengue-like symptoms. The prevalence of DENV decreased from 2014 to 2015 with the emergence of CHIKV. Genetic analyses indicate that there is movement of the DENV between Ecuador and neighboring countries, highlighting the importance of sentinel surveillance sites, such as Machala, in border regions. The rapid surveillance methods developed in this study could be applied to estimate the burden of other underreported febrile diseases, allowing the public health sector to more effectively and equitably conduct disease control interventions.

### Burden of DENV infection

Over the two years of the study, one third of associates had acute or recent DENV infections, a higher prevalence than findings from similar studies in Asia. In Vietnam, studies found 18% DENV prevalence in 100 meter clusters around initiate index cases, using PCR, NS1 ELISA, or serology.^21^ In Thailand, cluster DENV prevalence ranged from 10.1% to 14.3% using PCR or serology.^22,23^ One of possible explanations for the higher cluster prevalence in this study is the use of the NS1 rapid test. Prior studies that evaluated the Panbio Dengue Early Rapid test (used in this study) found that using antigen (NS1) and antibody (IgM, IgG) tests together increased the sensitivity of DENV diagnostics (93% sensitivity), and expanded the window of detection of infection.^52^ We found that the prevalence of DENV infections in clusters varied by DENV serotype (DENV1: 20.0%; DENV2: 36.6%). The higher cluster prevalence for DENV2 is consistent with prior studies that found greater infection rates for DENV2 compared to DENV1.^53^ The cause of the difference in infection rates between the two serotypes is not understood. Potential factors that could be involved include the local epidemiology, serotype subtype, weather, and previous exposure history of the population.^54^^‒^^56^

Using this active cluster surveillance protocol, we were able to effectively detect additional DENV infections in the community, particularly in 2014, when there was a higher burden of disease. For every initiate index case captured by passive surveillance, we captured approximately three associates with acute or recent (AR) DENV infections, of which two associates had symptomatic acute or recent (SAR) DENV infections. Interestingly, we found that the number of DENV infections per cluster was higher in 2014 than 2015, suggesting a higher force of DENV infection in 2014, when all four DENV serotypes were circulating, prior to the emergence of CHIKV. We temper this with caution, however, as our cluster sample size was smaller in 2015 (n=12) than 2014 (n=32).

In Latin America, enhanced surveillance studies that have reported DENV infection rates relative to passive surveillance infection rates include pediatric and adult cohorts, door-to-door community based surveillance studies, use of sentinel clinics, and enhanced laboratory diagnostic studies. To our knowledge, most cluster-based DENV surveillance studies with a similar design (*e.g.*, spatially restricted around the index home) have been conducted in Asian countries. Estimates of the burden of disease from active surveillance studies in Latin America vary widely depending on the study design, the effectiveness of passive surveillance, and the traits of the local population (*e.g.*, past exposure to DENV serotypes). In a pediatric cohort in Nicaragua, investigators detected 21.3 times more DENV infections than were reported to the national surveillance system.^57^ A study in Peru compared passive surveillance of DENV to a cohort study and sentinel clinic surveillance, and found five times more DENV infections in the cohort and 19 times more DENV infections through sentinel clinic surveillance.^25^ They found that both sentinel and cohort surveillance methods detected an increase in DENV infections more rapidly than passive surveillance methods. In Puerto Rico, laboratory enhanced surveillance resulted in three times more DENV infections registered than passive surveillance methods.^27^

One of the limitations of this study was that we surveyed the nearest neighbors of the initiate index case, which are not necessarily representative of the total population residing within 200 meters. We did not collect information on those who were not willing to participate in the study. Also, people may have been more willing to participate in the study if they or someone in their household was ill. This could potentially result in a higher estimate of the number of additional DENV infections in clusters compared to the general population. Future studies could survey a greater number of households located randomly within the 200-meter radius for a more accurate measure of disease prevalence and could assess DENV negative clusters as controls. Additionally, this study was limited to five clinical sites operated by the MoH that were willing and able to support the study. Testing for CHIKV and ZIKV was limited to PCR, and did not include serological testing.

### Burden of CHIKV and other febrile illness

In 2015, we found that 43.1% of clinically diagnosed (suspected) DENV infections were actually positive for CHIKV, higher than the proportion of laboratory-confirmed DENV infections. We identified six index cases and one associate with evidence of both acute CHIKV and acute or recent DENV infections in 2015 (11.5% of CHIKV infections). There were also 96 individuals with undiagnosed febrile illness (non-DENV, non-CHIKV, non-ZIKV). The burden of CHIKV is likely higher than reported here, since we only tested for acute infections. This highlights the difficulties of differential diagnosis in areas where DENV, CHIKV, ZIKV, and other febrile illnesses are co-circulating. These data also suggest that the large increase in DENV cases in 2015 in Ecuador (44,104 cases in 2015 versus 14,312 cases on average from 2010 to 2014)^11^ could be the result of CHIKV and other circulating febrile pathogens.

We did not detect ZIKV during the study period, consistent with MoH reports, which indicated that ZIKV circulated for the first time in Machala in February 2016. Although surveillance efforts were not focused specifically on clinical ZIKV infections, we suspect that the study would have detected some ZIKV infections if they were present in Machala due to the overlapping clinical presentations of DENV and ZIKV infections. However, recent studies indicate that urine and whole blood may be better suited to detect ZIKV, limiting our ability to detect ZIKV in serum samples by RT-PCR.^58,59^

### Clinical characteristics of DENV and CHIKV infections

In general, the symptoms that were observed with acute DENV infections in this study are consistent with other reports.^60^^‒^^66^ As in other studies, we found that secondary DENV infections were more severe; nine out of ten hospitalized individuals with DENV infections had secondary infections (Supplementary Table 4).^24,65,67^ From 2014 to 2015, we observed a shift from DENV2 to DENV1, and a shift from secondary to primary DENV infections. As expected, associates with acute DENV infections in 2015 were younger (mean=19.6 years of age) than in 2014 (mean=25.2 years of age), although the differences were not significantly different (Table 4). The clinical characteristics associated with DENV infections can vary over time and space due to both differences in the dominant serotypes in circulation^68,69^ and the ratio of primary to secondary infections.^24,65,67^

People infected with CHIKV versus DENV were older on average, consistent with the disease being newly introduced into the population. MoH reports indicated that the highest burden of CHIKV in Machala was among adults aged 20 to 49. We found that muscle and joint pain and rash were more commonly reported by people with CHIKV infections than those with DENV, which supports findings from prior studies.^62,66^

The ratio of symptomatic:inapparent (S:I) DENV infections in associates was 1:0.47 (2.13), which is within the upper range of prior estimates from DENV endemic regions. By defining symptomatic as any dengue-like symptom, rather than only fever, we captured a broad spectrum of DENV illness. Prior studies suggest that the S:I ratio for DENV infections can vary widely, possibly depending on the immune response to prior exposure to DENV serotypes, the serotypes (and subtypes) in circulation, and genetic factors.^23,24,31,33,69,70^ A one-year contact cluster study from Peru reported an S:I ratio of 1:4.56 (0.22).^31^ A four-year pediatric cohort study from Nicaragua reported S:I ratios ranging from 1:18.4 (0.05) to 1:3.0 (0.33).^69^ S:I ratios from a five-year school cohort study in Thailand ranged from greater than 4 to 0, depending on the year and school.^33,70^ A two-year school cohort and cluster study from Thailand reported an overall S:I ratio of 1:1 (1.0),^23^ and a one-year cluster surveillance study from Thailand reported 1:0.2 (5.0) for primary infections and 1:0.4 (2.5) for secondary infections.^23^ Differences may also be due to the profile of the study population (*e.g.*, adult versus pediatric) and how investigators defined symptomatic.

Despite the high proportion of associates with symptomatic acute DENV infections, few (5.9%) had sought medical care. In prior studies in Machala, community members and healthcare professionals indicated that there was low health care seeking behavior in certain populations, such as working men in the urban periphery, and self-medicating was common practice.^18,71^ Another explanation is that our definition of symptomatic DENV infections included mildly symptomatic infections that did not require medical attention. These findings highlight the importance of active surveillance protocols that capture inapparent infections and infections in demographic groups who are less likely to seek health care or who have limited access to health care.

### Phylogenetic analysis

Phylogenetic analyses of DENV1 showed Ecuadorian samples falling into two distinct clusters, sharing a common ancestor with viruses from Colombia in one cluster and a common ancestor with viruses from Venezuela in the other cluster. These well-separated clusters indicate at least two distinct introductions of DENV1 into Ecuador. Given the early sampling of Venezuelan and Colombian genomes (between 2004 and 2008), and given that recent DENV1 full genome samples from Peru are not available, we cannot exclude with certainty the role that Peru may have played in the DENV1 introductions into Ecuador. However, the results suggest a close genetic relationship of viruses circulating in Venezuela and Colombia and support the notion of commonly occurring DENV1 flow between the countries. Similar to DENV1, DENV2 genomes from Ecuador were most closely related to genomes from Venezuela and Colombia. However, unlike DENV1, DENV2 genomes from Ecuador made up a single monophyletic clade separated from the rest of the South American taxa with high support. This indicates a single introduction and subsequent spread of this virus in Ecuador without further DENV2 introductions and mixing from other regions. Even though older sequences from Peru clustered further away from genomes sampled in Ecuador, Venezuela, and Colombia, suggesting they did not play a role in the current DENV2 epidemic in Ecuador, the lack of recent full genomes from Peru prevent us from determining the involvement of Peru in the observed DENV2 spread in Ecuador. The unavailability of recent full genomes from countries surrounding Ecuador was most evident in DENV4, where the exact placement of the only Ecuadorian genome in the tree could not be determined due to low node support. Nevertheless, the results suggested a close relationship between DENV4 in Ecuador, Venezuela, Colombia and Brazil. It is important to note that samples from Peru were missing here as well, and that there is a possibility this country was also involved in the circulation of DENV4 in this region. Thus, our results suggest frequent flow of DENV between Ecuador and surrounding countries, including introduction and re-introduction of different serotypes and different lineages of the same serotype. In addition, our results show the importance of continuous surveillance, including genetic sequencing efforts. If available, virus full genomes from these countries would allow for more accurate analysis of the patterns of DENV movement and spread in this region.

### Public health implications

This study provides one of the most thorough descriptions of DENV and CHIKV infections in this region, and contributes to a long-term collaboration with the MoH and other governmental and academic partners to strengthen infectious disease surveillance in southern coastal Ecuador, a strategic area to monitor endemic and emerging pathogens. The collaboration has been successful due to a shared vision for integrated active surveillance that includes the virus-vector-host, climate and other social-ecological drivers;^20,32^ ongoing training of physicians, researchers and students; and improvement of local diagnostic and research infrastructure.

Enhanced surveillance studies, such as this, provide high-resolution spatiotemporal data on the distribution of symptomatic and inapparent infections across the population. This is especially important in places and in subgroups with low healthcare seeking behavior, which result in underreporting and continued disease transmission.^18,71^ Enhanced surveillance systems have been shown to detect an increase in infections earlier than passive surveillance systems,^25^ providing a warning of an escalating outbreak. These data are currently being used to parameterize and calibrate local epidemic forecast models.^72,73^ These data also allow the public health sector to more accurately estimate the social and economic cost of the disease, allowing for informed decision making regarding the allocation of scarce resources for current and future interventions, such as vector control, community mobilization, and vaccines.^74,75^ The age-stratified prevalence data generated through this study design provides important information for the design of future vaccine trials and vaccination campaigns.

Genetic and phylogenetic analyses provided additional information about virus movement and introductions into Ecuador. Determining sources of viral origin and most common pathways of spread provides important information about the dynamics of the epidemic that can aid in development of coordinated regional public health surveillance and control efforts, especially across Andean countries. Prior studies from the Ecuador-Peru border region highlight the importance of binational public health sector collaborations to effectively control mosquito-borne diseases.^76^ In addition, frequent movement of dengue between Ecuador and neighboring countries highlighted the importance of sentinel surveillance sites, such as Machala, in border regions.

## Acknowledgements

This project was possible thanks to support from colleagues from the Ministry of Health, the National Institute of Meteorology and Hydrology, the National Secretary of Higher Education, Science, Technology, and Innovation (SENESCYT) of Ecuador and community members from Machala, Ecuador. We thank our local field team and coordinators for their dedication and perseverance: Jefferson Adrian, Victor Arteaga, Jose Cueva, Reagan Deming, Carlos Enriquez, Prissila Fernandez, Froilan Heras, Naveed Heydari, Jesse Krisher, Lyndsay Krisher, Elizabeth McMahon, Eunice Ordoñez, and Tania Ordoñez. Many thanks to Rosemary Rochford, Lisa Ware, Holly Chanatry, David Amberg and Marti Benedict for supporting the development of the research platform with partners in Ecuador. We also thank Danielle Safaty and Laura Sorenson in the Center for Global Health and Translational Science at SUNY Upstate Medical University for technical support in sample preparation, RT-PCR analysis, and data compilation. We thank Dr. Renato Leon for supporting the development of the entomology protocol, and Ing. Raul Mejia and Dr. Angel Muñoz for supporting climate surveillance. Thank you to Dr. Butsaya Thaisomboonsuk PhD and Dr. Louis Macareo MD, JD from AFRIMS for sharing surveillance and diagnostic protocols. Thank you to Clinical Research Management (CRM) for supporting surveillance activities in 2016 and 2017.

## Disclaimer

Material has been reviewed by the Walter Reed Army Institute of Research. There is no objection to its presentation and/or publication. The opinions or assertions contained herein are the private views of the author, and are not to be construed as official, or as reflecting the views of the Department of the Army, or the Department of Defense.

## Current addresses of co-authors

Sadie J. Ryan: Department of Geography, University of Florida, Gainesville, FL, USA

Aileen Kenneson: U.S. Centers for Disease Control, Atlanta, GA, USA

Timothy P. Endy, Christine A. King, and Arturo Barbachano-Guerrero: Department of Microbiology & Immunology, SUNY Upstate Medical University, Syracuse, NY, USA

Mark Polhemus, Cinthya Cueva, Christina D. Lupone and Mark Abbott: Center for Global Health & Translational Sciences, SUNY Upstate Medical University, Syracuse, NY, USA

Efraín Beltrán-Ayala: Department of Medicine, Universidad

Técnica de Machala, Machala, El Oro Province, Ecuador

Mercy J. Borbor-Cordova and Washington B. Cárdenas: Department of Marine Engineering, oceanic and biological sciences, and natural resources. Escuela Superior Politecnica del Litoral (ESPOL), Guayaquil, Ecuador

Richard G. Jarman and Irina Maljkovic Berry: Viral Diseases Branch, Walter Reed Army Institute of Research (WRAIR), Silver Springs, MD, USA

Saurabh Mehta and Julia L. Finkelstein: Division of Nutritional Sciences, Cornell University, Ithaca, NY, USA

Mercy Silva: Ministry of Health, Machala, El Oro, Ecuador

---

**Table 5.** Demographics and symptoms associated with acute DENV infections versus acute CHIKV infections in index cases. Index cases with acute DENV infections were significantly younger and more likely to report anorexia and nausea, vomiting, and abdominal pain (p<0.05). Index cases with CHIKV were more likely to be female, were older, and more likely to report muscle/joint pain (p<0.05). One individual with a DENV and CHIKV co-infection was excluded.

| Characteristics | Acute DENV<br>N = 98 | Acute CHIKV<br>N = 52 | p-value |
| --- | --- | --- | --- |
| Age in years, mean (SD) | 20.2 (15.0) | 35.8 (19.4) | **<0.0001** |
| Gender, % female | 41/98 (41.8%) | 35/52 (67.3%) | **0.005** |
| Temperature > 38°C | 26/98 (26.5%) | 6/51 (11.8%) | 0.06 |
| Hospitalized | 20/98 (20.4%) | 5/52 (9.6%) | 0.14 |
| Symptoms in prior 7 days |  |  |  |
| Fever | 97/98 (99.0%) | 50/52 (96.2%) | 0.57 |
| Headache | 80/97 (82.5%) | 37/51 (72.5%) | 0.23 |
| Anorexia and nausea | 64/98 (65.3%) | 19/52 (36.5%) | **0.001** |
| Muscle/joint pain | 75/97 (77.3%) | 50/52 (96.2%) | **0.006** |
| Rash | 16/97 (16.5%) | 18/52 (34.6%) | 0.05 |
| Bleeding | 8/98 (8.2%) | 2/52 (3.8%) | 0.51 |
| Vomiting | 46/98 (46.9%) | 12/52 (23.1%) | **0.007** |
| Drowsiness/lethargy | 82/98 (93.9%) | 46/52 (88.5%) | 0.58 |
| Abdominal pain | 62/97 (63.9%) | 19/52 (36.5%) | **0.002** |
| Diarrhea | 27/98 (27.6%) | 16/52 (30.8%) | 0.82 |
| Retro-orbital pain | 67/98 (68.4%) | 35/51 (68.6%) | 1 |

**Supplementary Table 1.** The prevalence of dengue-like symptoms in associates with acute DENV infections. Dengue-like symptoms include all symptoms listed below. Symptoms are presented from most to least prevalent.

| Symptoms | N=50 | Prevalence |
| --- | --- | --- |
| Any dengue-like symptom | 34 | 68% |
| Temperature > 38°C | 2 | 4% |
| Symptoms in prior 7 days |  |  |
| Headache | 16 | 32% |
| Drowsiness/lethargy | 12 | 24% |
| Fever | 11 | 22% |
| Muscle/joint pain | 11 | 22% |
| Retro-orbital pain | 11 | 22% |
| Abdominal pain | 9 | 18% |
| Rash | 9 | 18% |
| Anorexia and nausea | 5 | 10% |
| Diarrhea | 3 | 6% |
| Vomiting | 2 | 4% |
| Bleeding | 1 | 2% |

**Supplementary Table 2.**
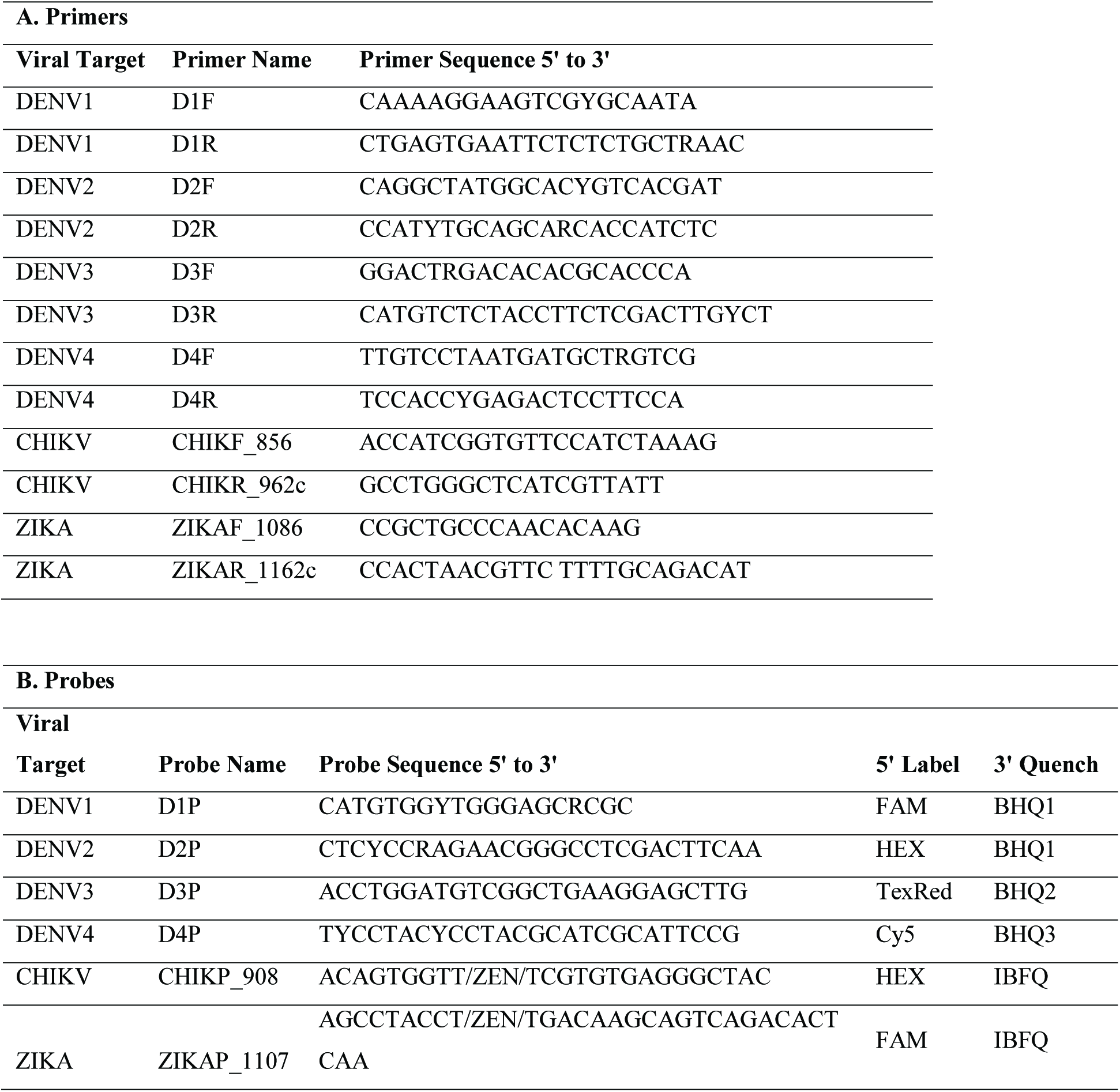
(A) Primers and (b) probes used for RT-PCR diagnostics of DENV, CHIKV, and ZIKV.

**Supplementary Table 3.**
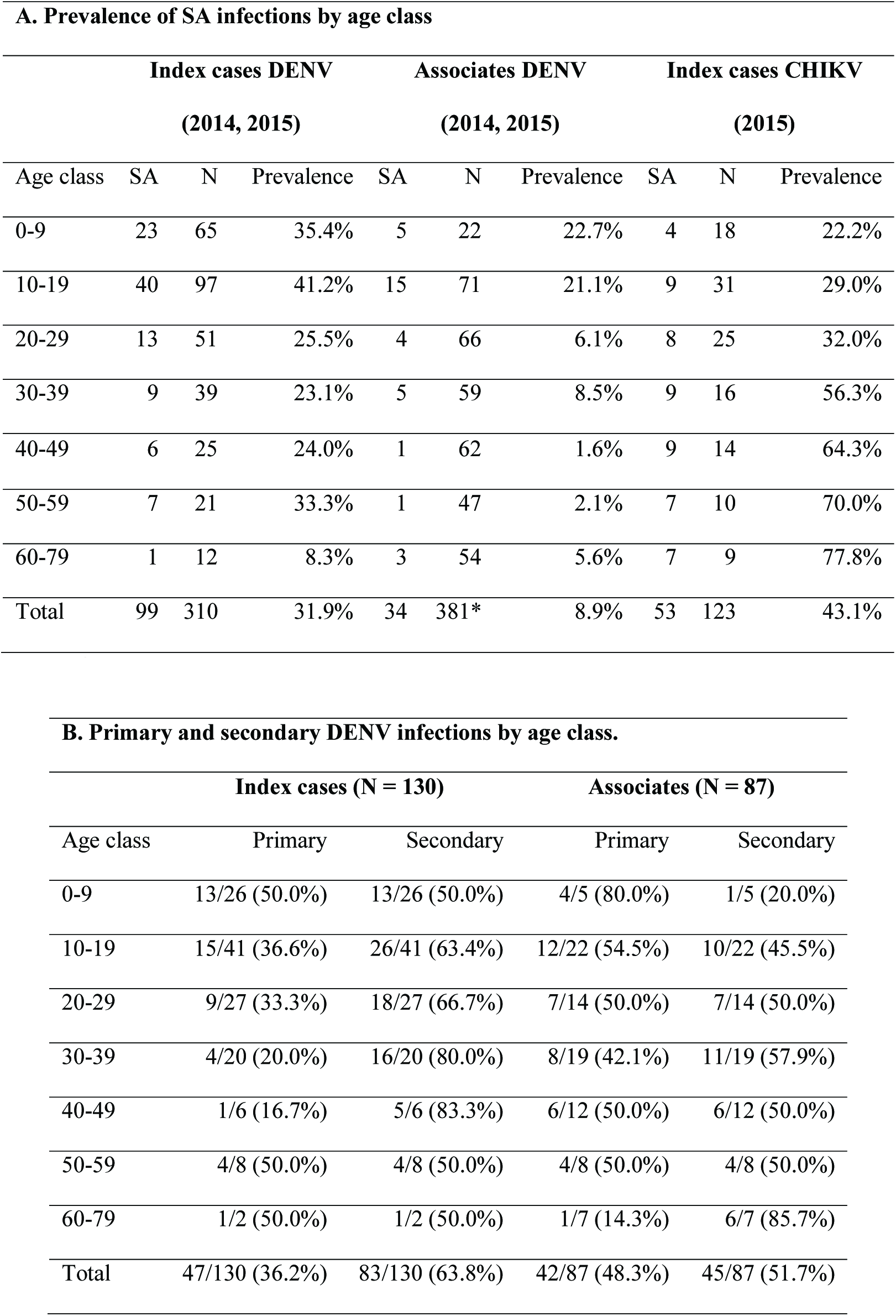
The prevalence of symptomatic acute (SA) infections and serology by age class. Data were used to generate Figure 4. (A) Index cases and associates with symptomatic acute (SA) DENV or CHIKV infections, as a proportion of all individuals from the age class who were recruited into the study (N). For DENV, data are combined for 2014 and 2015. For CHIKV, data are shown only for 2015. There were no associates with SA CHIKV infections. (B) The proportion of primary and secondary DENV infections per age class for index cases and associates with valid serology and acute or recent DENV infections in 2014 and 2015 combined.

*3 associates were missing age information.

**Supplementary Table 4.**
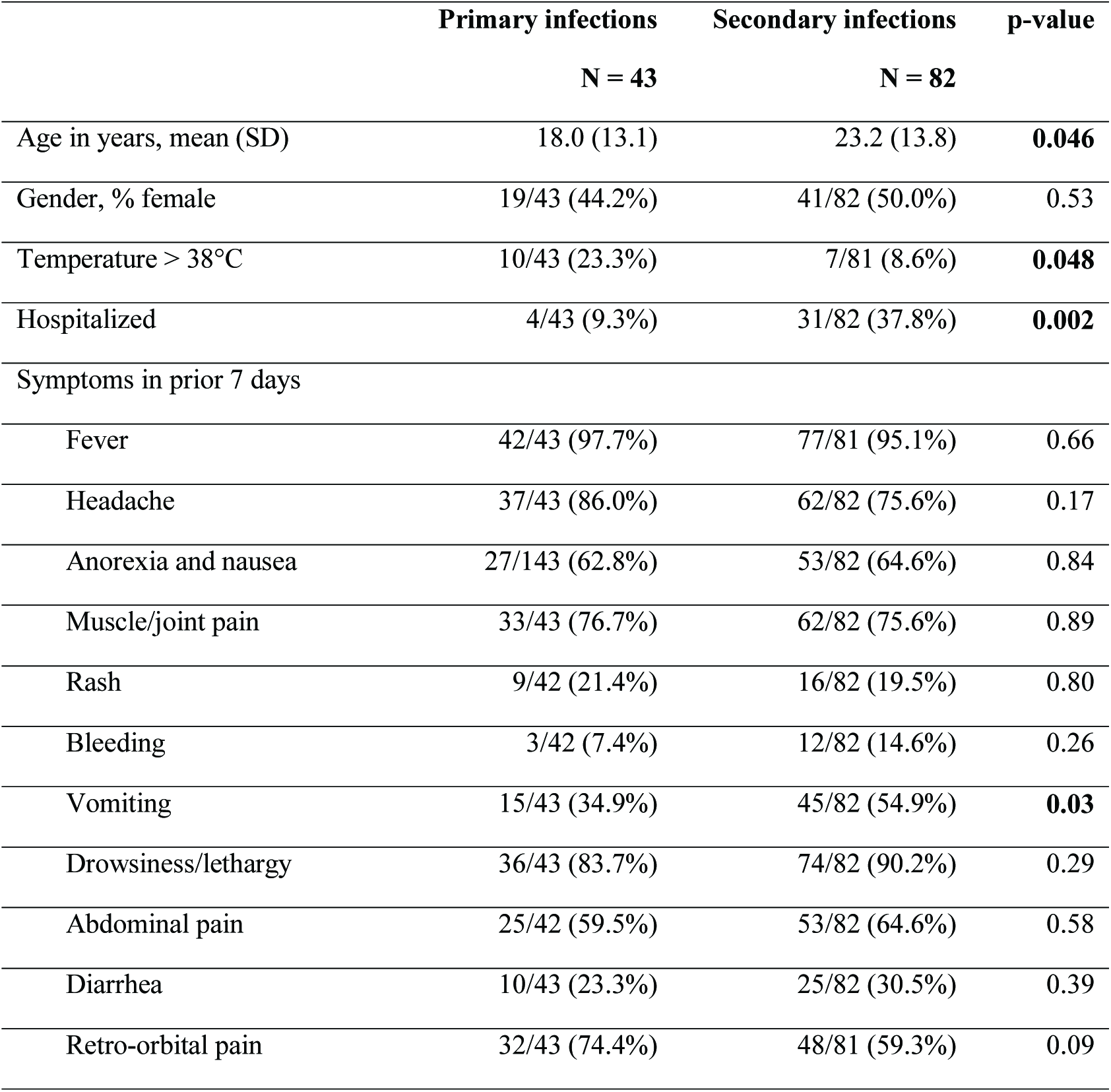
Demographics and symptoms associated with primary versus secondary DENV infections in index cases that had acute or recent DENV infections. Index cases with secondary DENV infections were significantly older, were less likely to have a fever, and were more likely to report vomiting (p<0.05). Hospitalized cases were more likely to have secondary infections. Index cases with DENV and CHIKV co-infections were excluded (4 primary infections, 1 secondary infection).

**Supplementary Table 5.** Demographics and symptoms associated with DENV1 versus DENV2 infections in index cases. Index cases with DENV1 infections were significantly younger than those with DENV2 infections (p<0.05). For all other measures, there were no significant differences (p>0.05). One index case with a DENV and CHIKV co-infection was excluded.

|  | **DENV1** | **DENV2** | **p-value** |
| --- | --- | --- | --- |
|  | **N = 18** | **N = 51** |  |
| Age in years, mean (SD) | 14.7 (10.5) | 25.2 (16.2) | **0.01** |
| Gender, % female | 9/18 (50.0%) | 21/51 (41.2%) | 0.71 |
| Temperature > 38°C | 8/18 (44.4%) | 15/51 (29.4%) | 0.38 |
| Hospitalized | 5/18 (27.8%) | 7/51 (13.7%) | 0.32 |
| Symptoms in prior 7 days |  |  |  |
| Fever | 18/18 (100%) | 49/51 (96.1%) | 0.97 |
| Headache | 17/18 (94.4%) | 43/51 (84.3%) | 0.49 |
| Anorexia and nausea | 14/18 (77.8%) | 32/51 (62.8%) | 0.38 |
| Muscle/joint pain | 12/18 (66.7%) | 43/51 (84.3%) | 0.21 |
| Rash | 2/17 (11.8%) | 8/51 (15.7%) | 1.00 |
| Bleeding | 3/18 (16.7%) | 2/51 (3.92%) | 0.21 |
| Vomiting | 9/18 (50.0%) | 26/51 (51.0%) | 1.00 |
| Drowsiness/lethargy | 16/18 (88.9%) | 44/51 (86.3%) | 1.00 |
| Abdominal pain | 13/18 (72.2%) | 31/51 (60.8%) | 0.56 |
| Diarrhea | 4/18 (22.2%) | 12/51 (23.5%) | 1.00 |
| Retro-orbital pain | 13/18 (72.2%) | 36/51 (70.6%) | 1.00 |

**Supplementary Table 6.**
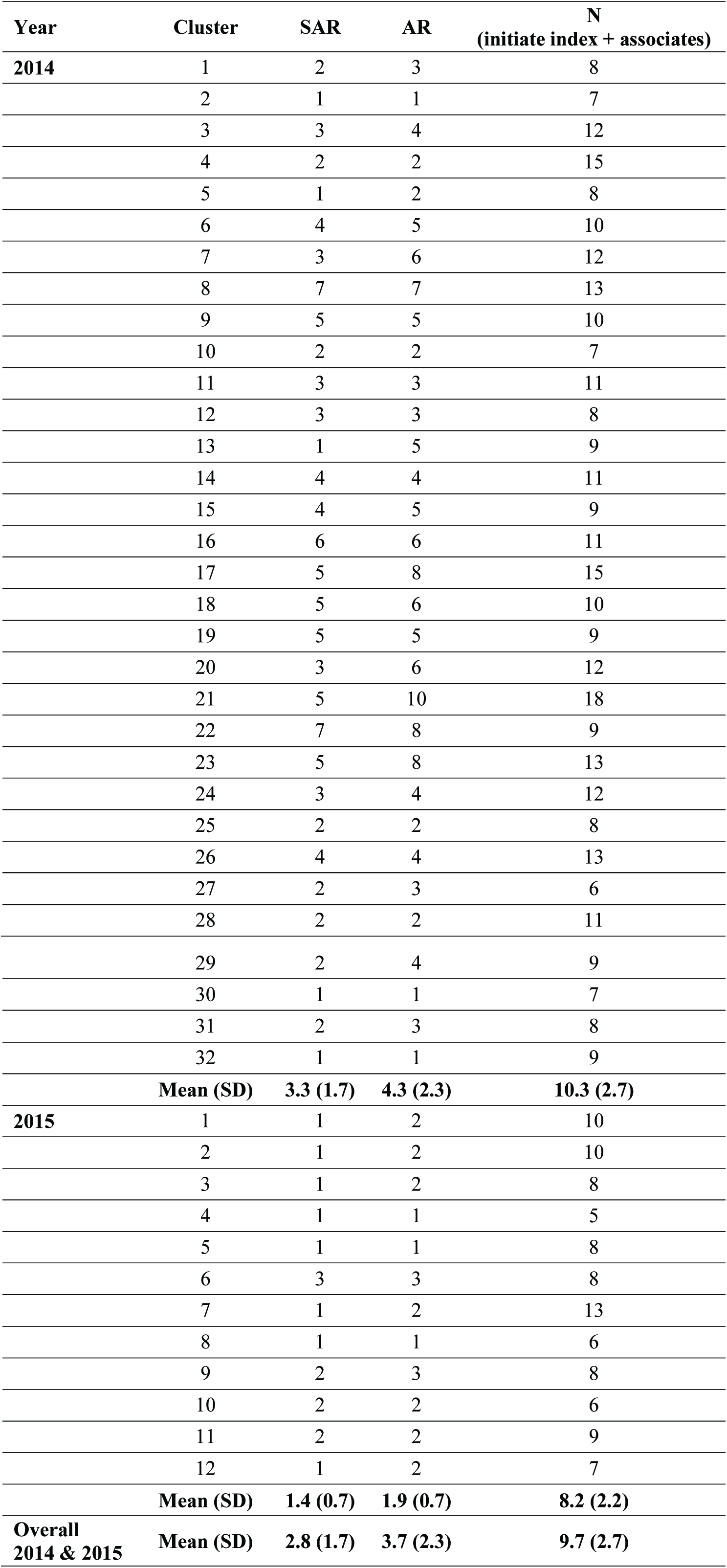
DENV infections per cluster. The numbers of symptomatic acute and recent (SAR) DENV infections, and acute and recent (AR) DENV infections per cluster, and the total number of people per cluster. Each cluster includes one initiate index case, which by definition was a SAR infection. Means and standard deviations (SD) for clusters are shown for each year and for both years combined. All measures were significantly greater in 2014 than in 2015 (p<0.05).

| Year | Cluster | SAR | AR | N<br>(initiate index + associates) |
| --- | --- | --- | --- | --- |
| 2014 | 1 | 2 | 3 | 8 |
|  | 2 | 1 | 1 | 7 |
|  | 3 | 3 | 4 | 12 |
|  | 4 | 2 | 2 | 15 |
|  | 5 | 1 | 2 | 8 |
|  | 6 | 4 | 5 | 10 |
|  | 7 | 3 | 6 | 12 |
|  | 8 | 7 | 7 | 13 |
|  | 9 | 5 | 5 | 10 |
|  | 10 | 2 | 2 | 7 |
|  | 11 | 3 | 3 | 11 |
|  | 12 | 3 | 3 | 8 |
|  | 13 | 1 | 5 | 9 |
|  | 14 | 4 | 4 | 11 |
|  | 15 | 4 | 5 | 9 |
|  | 16 | 6 | 6 | 11 |
|  | 17 | 5 | 8 | 15 |
|  | 18 | 5 | 6 | 10 |
|  | 19 | 5 | 5 | 9 |
|  | 20 | 3 | 6 | 12 |
|  | 21 | 5 | 10 | 18 |
|  | 22 | 7 | 8 | 9 |
|  | 23 | 5 | 8 | 13 |
|  | 24 | 3 | 4 | 12 |
|  | 25 | 2 | 2 | 8 |
|  | 26 | 4 | 4 | 13 |
|  | 27 | 2 | 3 | 6 |
|  | 28 | 2 | 2 | 11 |
|  | 29 | 2 | 4 | 9 |
|  | 30 | 1 | 1 | 7 |
|  | 31 | 2 | 3 | 8 |
|  | 32 | 1 | 1 | 9 |
|  | **Mean (SD)** | **3.3 (1.7)** | **4.3 (2.3)** | **10.3 (2.7)** |
| **2015** | 1 | 1 | 2 | 10 |
|  | 2 | 1 | 2 | 10 |
|  | 3 | 1 | 2 | 8 |
|  | 4 | 1 | 1 | 5 |
|  | 5 | 1 | 1 | 8 |
|  | 6 | 3 | 3 | 8 |
|  | 7 | 1 | 2 | 13 |
|  | 8 | 1 | 1 | 6 |
|  | 9 | 2 | 3 | 8 |
|  | 10 | 2 | 2 | 6 |
|  | 11 | 2 | 2 | 9 |
|  | 12 | 1 | 2 | 7 |
|  | **Mean (SD)** | **1.4 (0.7)** | **1.9 (0.7)** | **8.2 (2.2)** |
| **Overall**<br>**2014 & 2015** | **Mean (SD)** | **2.8 (1.7)** | **3.7 (2.3)** | **9.7 (2.7)** |

